# Mammary epithelial cells have lineage-restricted metabolic identities

**DOI:** 10.1101/798173

**Authors:** Mathepan Mahendralingam, Kazeera Aliar, Alison Elisabeth Casey, Davide Pellacani, Hyeyeon Kim, Vladimir Ignatchenko, Mar Garcia Valero, Luis Palomero, Ankit Sinha, Vid Stambolic, Mina Alam, Aaron Schimmer, Hal Berman, Miquel Angel Pujana, Connie Eaves, Thomas Kislinger, Rama Khokha

## Abstract

Cancer metabolism adapts the metabolic network of its tissue-of-origin. However, breast cancer is not a disease of a singular origin. Multiple epithelial populations serve as the culprit cell-of-origin for specific breast cancer subtypes, yet knowledge surrounding the metabolic network of normal mammary epithelial cells is limited. Here, we show that mammary populations have cell type-specific metabolic programs. Primary human breast cell proteomes of basal, luminal progenitor, and mature luminal populations revealed their unique enrichment of metabolic proteins. Luminal progenitors had higher abundance of electron transport chain subunits and capacity for oxidative phosphorylation, whereas basal cells were more glycolytic. Targeting oxidative phosphorylation and glycolysis with inhibitors exposed distinct metabolic vulnerabilities of the mammary lineages. Computational analysis indicated that breast cancer subtypes retain metabolic features of their putative cell-of-origin. Lineage-restricted metabolic identities of normal mammary cells partly explain breast cancer metabolic heterogeneity and rationalize targeting subtype-specific metabolic vulnerabilities to advance breast cancer therapy.

## INTRODUCTION

Molecular classification of breast cancers using the PAM50 classifier has identified 5 main patient groups (Luminal A, Luminal B, HER2, Claudin-low, Basal-like) with distinct transcriptional programs, survival outcomes and susceptibilities to anti-cancer regimens^1–3^. Metabolomics on primary tumors report distinct metabolic phenotypes for each of the breast cancer subtypes^4–8^. Investigation of subtype-specific metabolic features have focused on the effects of estrogen receptor^9^, *HER2* amplification^10^ and/or driver mutations (*Tp53*, *Pik3ca*)^11^. However, several of these markers and mutations are shared amongst breast cancer subtypes and can only partly explain the metabolic heterogeneity^12^. Tissue-of-origin has emerged as an important intrinsic determinant of cellular metabolism^13^. This is based on studies showing cancers use the metabolic network of their normal counterpart as a backbone for aberrant proliferation^14–16^. This poses a challenge in the context of breast cancer as there are multiple cell(s)-of-origin^17–19^, each postulated to give rise to a specific breast cancer subtype. Whether individual precursor cells from a single tissue have intrinsic differences in their metabolism remains unknown.

The mammary gland is composed of two epithelial lineages, the basal and luminal lineages. Milk-producing luminal cells and contractile basal cells operate in unison to carry out the overall function of the breast^20^. The luminal lineage can be segregated into luminal progenitors and mature luminal (more differentiated) populations, whereas markers to segregate subpopulations within the basal lineage have not been conclusively defined (Figure 1A). Each of these three normal mammary epithelial cell (MEC) types serve as the putative cell-of-origin for distinct breast cancer subtypes. Expression analyses have projected that basal cells give rise to the Claudin-low subtype, mature luminal cells to Luminal A & B and luminal progenitors transform to the aggressive Basal-like subtype^21^. Mouse models with lineage-specific promoters also support the observation that the same mutational event results in different breast cancers depending upon the cell-of-origin^22–24^. Transcriptomic and epigenomic profiling of both human and mouse normal MECs have revealed lineage-specific regulatory networks^25–27^. With respect to metabolism, fetal mammary stem cells had high transcript levels of glycolysis enzymes^28^ and luminal progenitors were shown to have a greater capacity to handle reactive oxygen species (ROS) than basal cells^29^. Nevertheless, the metabolic networks of normal mammary cell types have yet to be resolved and whether breast cancer subtypes retain these metabolic features from their distinct cells-of-origin remains unknown.

**Figure 1:**
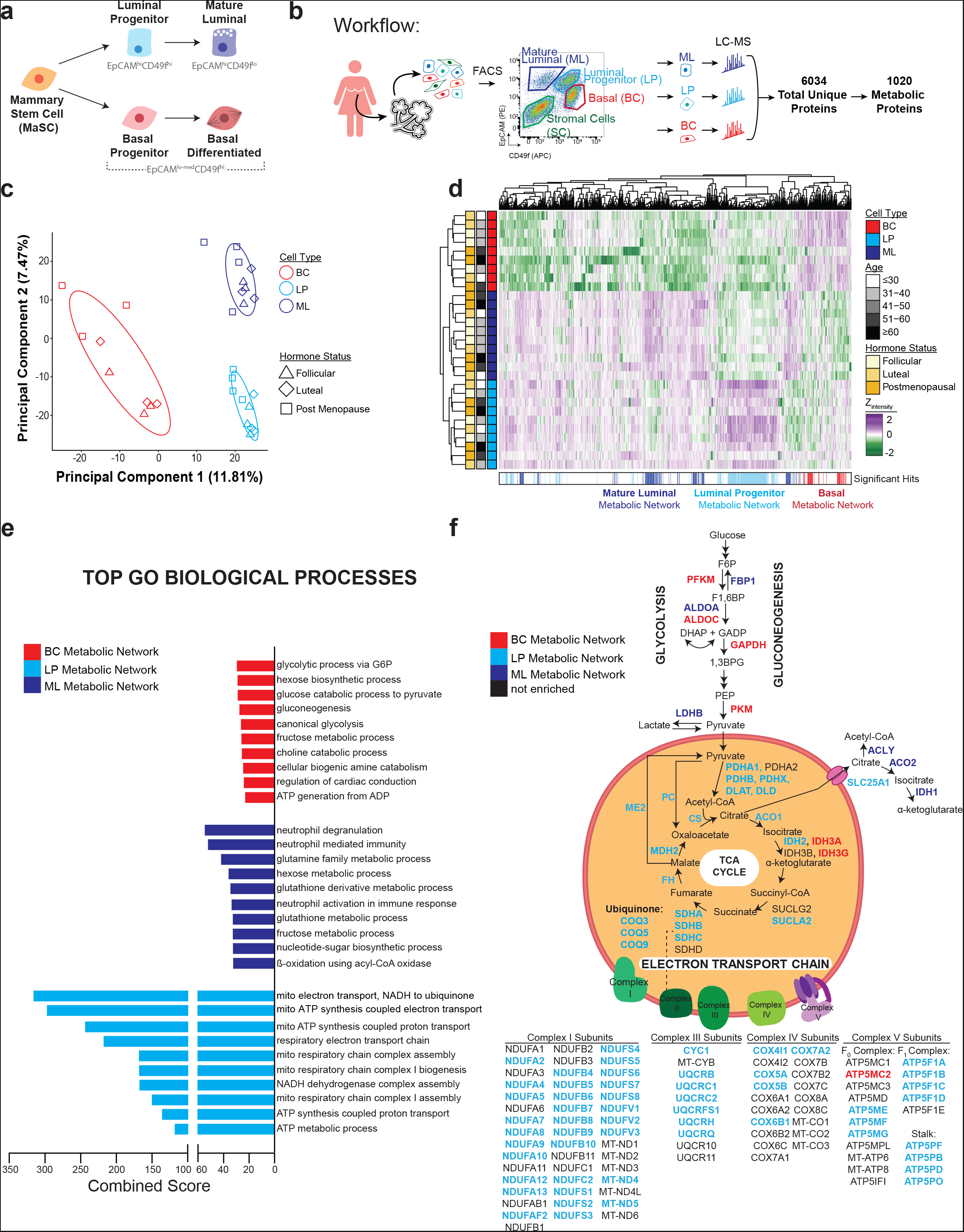
Proteomics illustrate distinct metabolic networks of human MECs. A. Mammary epithelial cell (MEC) hierarchy depicting the basal and luminal lineages and cell surface markers used to FACS-purify basal, luminal progenitor and mature luminal cells. B. Schematic depicting workflow on how human breast samples (n=10) were processed to single cells, the FACS gating strategy used to segregate mature luminal (ML), luminal progenitor (LP) and basal cells (BC) populations. Purified fractions were then prepared for liquid chromatography-mass spectrometry (LC-MS). Proteomics yielded 6034 uniquely detected proteins, whose abundance was corrected for batch effects, and missing values were imputed prior to downstream analyses. Total proteomes were filtered down to 1020 metabolic proteins. C. Principal component analysis of total proteome from human BC, LP and ML. Dot colour represents a mammary cell type, dot-shape represents hormone status (follicular, luteal or postmenopause) and ellipses represents clusters of sample types. D. Heatmap showing unsupervised hierarchical clustering and enrichment of the 1020 metabolic proteins in human MECs. Patient covariates (cell type, hormone status, age) are shown in the bars aligning the heatmap. Each line found in the “Significant Hits” bar is a metabolic protein whose expression was significantly enriched in only one cell type using a one-way ANOVA in conjunction with Tukey’s test (p<0.05) and colour-coded for that cell type. E. Bar graphs summarize the top 10 most significant GO biological processes according to Enrichr for each cell type’s metabolic network. Enrichr calculates a combined score, a metric used to find the best terms. F. Metabolic proteins participating in glycolysis, TCA cycle and ETC found in our proteomes are illustrated. Proteins that were significantly enriched in MEC-specific metabolic network appear in bold and are colour-coded to signify cell type.

Here, we uncover the distinct metabolic identities of the three normal mammary epithelial cell types by using a combination of proteomics, characterization of the mitochondria and pharmacological inhibition. In addition, their distinct metabolic networks not only underlie the differential dependencies of mammary progenitors to metabolic inhibitors, but are also inherited by the specific breast cancer subtypes.

## RESULTS

### Proteomes of Human Mammary Cells Expose Differential Metabolic Protein Abundance

To discover protein distinctions of primary human MEC populations we generated their global proteomes. We performed mass spectrometry-based shotgun proteomics on equivalent numbers of FACS-purified basal (CD45‾CD31‾CD49f^hi^EpCAM^lo-med^; color-coded as red in all figures), luminal progenitor (CD45‾CD31‾CD49f^lo^EpCAM^med^; light blue), and mature luminal (CD45‾CD31‾CD49f^hi^EpCAM^lo^; dark blue) cells from 10 normal human breast samples obtained from reduction mammoplasties (Figure 1B, S1A). Our patient cohort represented diverse physiologies, covering a wide age range (28-67 years old) and sex hormone status (3 luteal, 3 follicular, 4 post-menopausal). We detected 6034 unique proteins (Figure 1B). Expression of known markers for each mammary cell type was accurately captured by our proteomics data (Figure S1B); higher abundance of Vimentin and ITGA6 (Integrin α6, CD49f) was seen in basal cells, higher KIT and ALDH1A3 levels in luminal progenitors, and higher GATA3, FOXA1 and KRT8/18 (Cytokeratin 8/18) in the mature luminal. Principal component analysis highlighted the distinct proteomes of mammary cells; the dominant clustering feature was mammary cell type with a minor segregation of post-menopausal samples within each cluster (Figure 1C). Out of the 6034 proteins, 5881 were detected in all three cell types (Figure S1C). MEC-specific proteomes separated into deciles based on median intensity were enriched for specific functional classes of proteins (Figure S1D and S1E)^30^. For instance, the GO biological process in the first decile for basal cells and luminal progenitors was translation of membrane proteins, whereas mature luminal demonstrated abundance for proteins with pleotropic functions that were annotated as neutrophil terms (Figure S1D and S1E).

A metabolic network is defined as the core set of metabolic proteins essential for the structure and function of a cell^13^. We first filtered the total proteomes for metabolic proteins using a curated list of 2753 metabolic enzymes, transporters and subunits^31^. One sixth (1020/6034) of our global mammary proteomic dataset was classified with this annotation (Figure 1B). Unsupervised hierarchical clustering of the metabolic proteome clustered based on mammary lineages (Figure 1D). To determine the metabolic network functioning within each MEC, we sought proteins that were significantly abundant in one population versus the other two (One-way ANOVA in conjunction with a Tukey’s test, P<0.05). The resulting metabolic networks for basal, mature luminal and luminal progenitor were composed of 45, 123, 179 metabolic proteins, respectively (significant hits bar in Figure 1D). Pathway analysis using Enrichr^32,33^ revealed unique GO Biological Processes enriched in each metabolic network as found in Figure 1E. We also constructed a global map of MEC metabolism (Figure S2) using a published template^34^ and one focused on glucose metabolism (Figure 1F), where we color-coded proteins to show their corresponding MEC-specificity.

Basal cells were enriched for GO terms relating to glycolysis (Figure 1E) and displayed an abundance of glycolytic enzymes (PFKM, ALDOC, GAPDH and PKM); PFKM and PKM perform two of three key irreversible phosphorylation events in glycolysis (Figure 1F). In the mature luminal metabolic network, we noted diverse pathways relating to neutrophil activity, glutamine and glutathione (Figure 1E). It was also enriched for enzymes in hexose and fructose metabolism such as FBP1, ALDOA and LDHB; LDHB diverts pyruvate from the TCA cycle by converting it to lactate (Figure 1F). It was striking that the top 10 pathways in luminal progenitors related to oxidative phosphorylation (OXPHOS; Figure 1E), demonstrating greater abundance of the majority of electron transport chain (ETC) subunits as well as nearly all enzymes in the TCA cycle (Figure 1F). MECs had isozyme-specific expression of IDH (mitochondrial IDH3 in basal cells, IDH2 in luminal progenitors and IDH1 in mature luminal), possibly due to different levels of (NAD(P)H) and demand of that in particular cell type^35^. Pyruvate generated from the carbons of glucose is considered a major contributor to the TCA cycle, however luminal progenitors did not display any enrichment of glycolytic enzymes. Interrogation of our MEC metabolism map (Figure S2) revealed numerous non-glycolytic mechanisms to generate TCA cycle intermediates in luminal progenitors. This was the only population to exhibit enrichment for enzymes involved in branched-chain amino acid catabolism (modifying isoleucine, leucine and valine into acetyl-CoA) and was also strongly enriched for proteins involved in β-oxidation (fatty acids into acetyl-CoA) (Figure S2). Luminal progenitors also had high level of PHGDH, a key enzyme in serine biosynthesis, shown to contribute ~50% of the anaplerotic flux into the TCA cycle^31^ (Figure S2). This engagement of diverse metabolic pathways that break down nutrients to feed the TCA cycle underscores the strong preference of OXPHOS in this cell type. Altogether, proteomes revealed that each mammary cell type has a distinct metabolic network, which may represent the core set of metabolic proteins necessary for its structure and function.

### Mitochondria Structure and Function is Mammary Cell Type-Specific

Next, we interrogated our published mouse mammary proteomic dataset^25^ derived from analogous MEC populations and found that metabolic proteomes clustered based on mammary cell types (Figure S3A and S3B), similar to human MECs (Figure 1D). Since human luminal progenitors were endowed with TCA cycle and ETC proteins, we utilized murine MECs cultured as a monolayer to examine their capacity to undergo OXPHOS as measured by the Seahorse bioanalyzer (Figure 2A and 2B). Specifically, we performed the standard mitochondrial stress test, which quantifies oxygen consumption rate (OCR), a readout for mitochondrial respiration, while exposing cells to inhibitors (Oligomycin, Antimycin A) or enhancers (FCCP) of this process. At baseline respiration, basal cells had the lowest level of OCR compared to either of the luminal populations (Figure 2B). Even with the addition of FCCP, which boosts OCR, basal cells had OCR levels comparable to or less than the baseline OCR of the two luminal cell types. Luminal progenitors and mature luminal cells had similar OCR profiles, except for maximal respiration, which was significantly higher in luminal progenitors (Figure 2B). Thus, mammary cell types have distinct capacities for mitochondrial respiration, with luminal progenitors showing the highest OXPHOS capacity.

**Figure 2:**
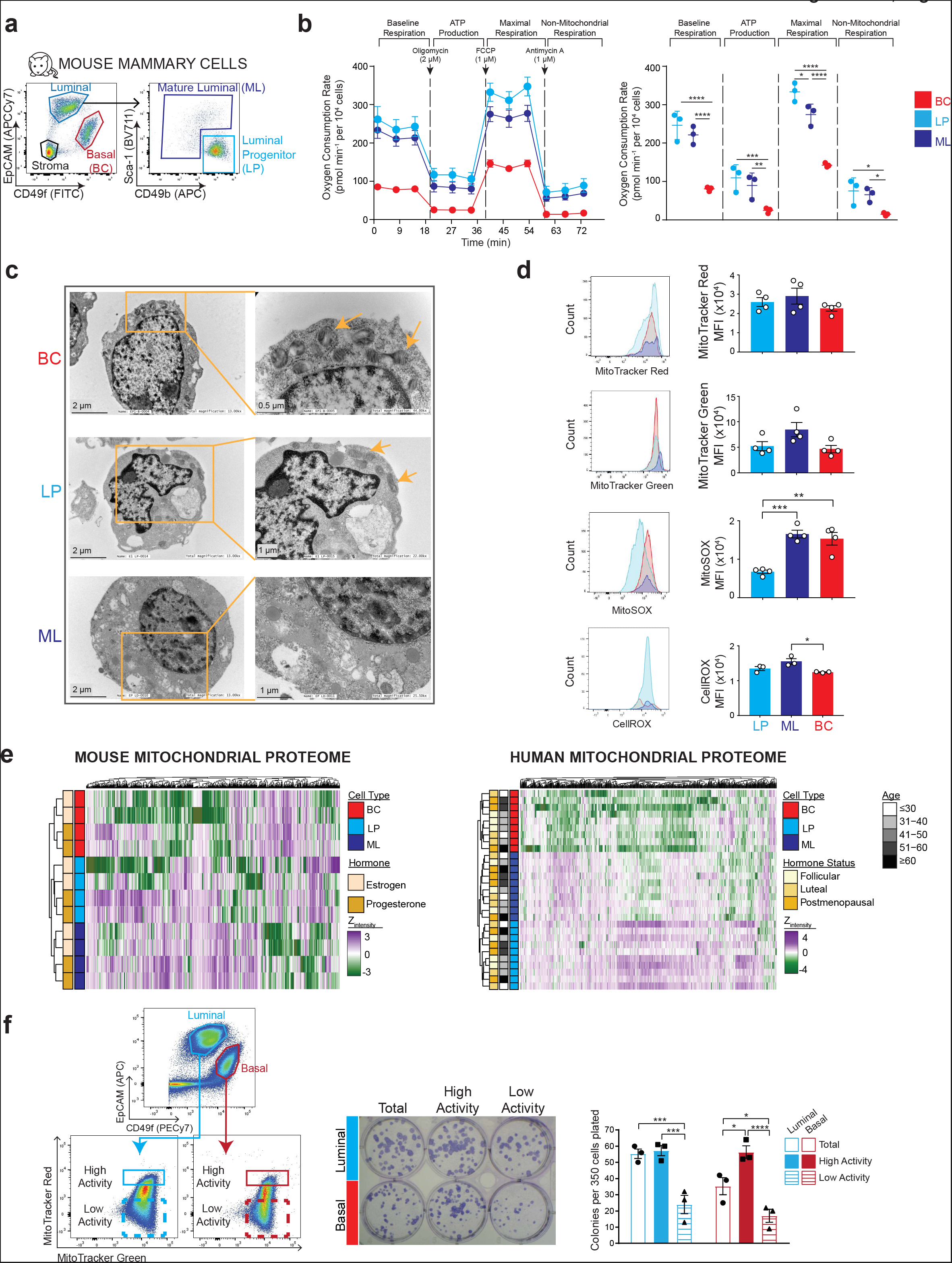
Mitochondrial structure and function varies with mammary lineage. A. FACS gating strategy to purify analogous mouse MEC populations. B. Oxygen consumption rate, determined by Seahorse Bioanalyzer, of mouse MECs at baseline, after exposure to Oligomycin (2 μM), FCCP (1 μM) and Antimycin A (1 μM). The left panel depicts the kinetic view of the data, which is quantified in the right panel (n = 3 mice; 4 technical replicates per n). All data are mean ± SEM. * P≤0.05; ** P≤0.01; *** P≤0.001; ****P≤0.0001. C. Representative transmission electron micrographs of FACS-sorted mammary cell pellets. Arrows indicate mitochondria. Magnifications are specified in each image. D. Flow plots and quantification of median fluorescent intensity (MFI) for MitoTracker Red (mitochondrial activity), MitoTracker Green (total mitochondria), MitoSOX (mitochondrial ROS) and CellROX (total ROS). Each dot represents a biological replicate (n=3-4 mice). E. Heatmap showing unsupervised hierarchical clustering and z-scores of mitochondrial protein abundance in mouse and human mammary proteomes with defined sex hormone status and patient characteristics. MitoCarta^40^, a curated list of mitochondrial proteins, was used to filter our total MEC-specific proteomes. The mouse proteome was obtained from a recent publication ^25^. F. Gating strategy used to sort basal and luminal cells with high and low mitochondrial activity (left) that were subsequently cultured in the colony forming cell (CFC) assay. Representative colony images and quantification of colonies formed from culturing luminal and basal cells with either high or low mitochondrial activity in CFC assay. Each dot represents a biological replicate (n=3 mice).

We next examined mitochondrial morphologies by transmission electron microscopy (TEM) of pelleted FACS purified mouse MEC (Figure 2C and S3C). Basal cells tended to have several small circular mitochondria with glossy cristae. This is similar to the morphology of mitochondria in hematopoietic and embryonic stem cells^36^. In contrast, luminal progenitors had long, tubular mitochondria with elaborate cristae. The larger size of the mitochondria and higher cristae density are thought to be efficient in supporting OXPHOS^37^, consistent with our data on OCR (Figure 2B). Mature luminal cells surprisingly had indiscernible mitochondria, possibly due to the shearing stress experienced by these larger cells during FACS. We performed intracellular flow cytometry to further characterize the mitochondria (Figure 2D), using MitoTracker Green (MTG; total level of mitochondria) and MitoTracker Red (MTR; mitochondrial activity). Both dyes showed no significant differences among MECs (Figure 2D). CellROX and MitoSOX measure cellular and mitochondrial reactive oxygen species (mROS), respectively. Although total cellular ROS showed minimal differences, mROS levels varied significantly (Figure 2D). Basal and mature luminal cells had equivalent high levels of mROS, whereas luminal progenitors had the least amount despite having high mitochondrial respiration (Figure 2D). These observations can be explained by the multiple antioxidant mechanisms previously reported in luminal progenitors but not in basal cells^29^.

The high mROS levels in basal cells were intriguing, as they did not have high OCR. This led us to ask whether the mitochondria had an alternative role in this population beyond bioenergetics. The non-ATP functions of the mitochondria are becoming more appreciated. For example, mitochondrial membrane potential has been linked to stem cell capacity^38,39^. We filtered our mouse and human MEC proteomes using MitoCarta, a curated list of mitochondrial proteins^40^, and observed cell type-based clusters by unsupervised hierarchical clustering (Figure 2E). Basal and luminal lineages are each enriched for their own progenitors. We segregated MECs based on high (MTG^hi^MTR^hi^) or low (MTG^hi^MTR^lo^) mitochondrial activity^41^, and enumerated luminal and basal progenitor capacity using the colony-forming cell (CFC) assay (Figure 2F). Cells with high mitochondrial activity had significantly greater CFC number than those with low mitochondrial activity in both mammary lineages (Figure 2E). Basal, but not luminal, cells with high mitochondrial activity also displayed enrichment of CFC capacity when compared to their total unfractionated control. The EpCAM^−^CD49f^hi^ basal cell compartment contains mammary stem cells, basal progenitors and differentiated cells. There is avid interest in teasing out new markers for progenitor-enriched basal subsets and our data show that mitochondrial activity may serve such a role. Overall, our findings also show that mitochondrial morphology and function varies with mammary cell type.

### Mammary Lineages Exhibit Differential Metabolic Vulnerabilities

We determined if metabolic distinctions of mammary lineages manifested as differential sensitivity to various metabolic drugs using the CFC assay (Figure 3A). To inhibit OXPHOS, we used complex-specific (Rotenone→Complex I; Atpenin A5→Complex II; Antimycin A→Complex III; Oligomycin→Complex V) and non-ETC inhibitors (Tigecycline→mitochondrial ribosomes; UK5099→Mitochondrial pyruvate carrier). Furthermore, inhibition of glycolysis was achieved at multiple levels (BAY-876→Glucose transporter 1; 2-Deoxy-D-glucose→Hexokinase; Galloflavin→Lactate dehydrogenase; Dichloroacetate→Pyruvate dehydrogenase kinase). Most metabolic inhibitors resulted in a potent dose-dependent reduction in progenitor capacity of both lineages, as enumerated by absolute CFCs (Figure S4A & S4B). Relative CFC counts allow comparison of the selective vulnerability of luminal and basal progenitors to metabolic inhibitors (Figure 3B and 3C). Analyses of the dependencies show that the two mammary lineages require specific ETC complexes for their progenitor capacity. We observed that inhibition of Complex I preferentially decreased luminal CFCs (Figure 3B) whereas basal CFCs were significantly more sensitive to Complex II or III inhibition. Complex V inhibition showed no selective effect (Figure 3B). Tigecycline treatment abrogated the progenitor capacity of basal over luminal CFCs (Figure 3B), supporting our earlier data that mitochondria strongly influence basal progenitor activity (Figure 2F). UK5099 prevents entry of pyruvate into mitochondria, but had a minimal effect on CFC capacity (Figure 3B), in line with our observation that luminal progenitors may not rely on cytosolic pyruvate (Figure 1E and S2). Pathway analyses of the basal metabolic network had highlighted glycolysis as the most significant term (Figure 1E). Our series of glycolytic drugs demonstrated that basal CFCs were far more sensitive than luminal CFCs to all 4 compounds (Figure 3C). Collectively, this set of experiments demonstrates the lineage-specific metabolic vulnerabilities of mammary cells.

**Figure 3:**
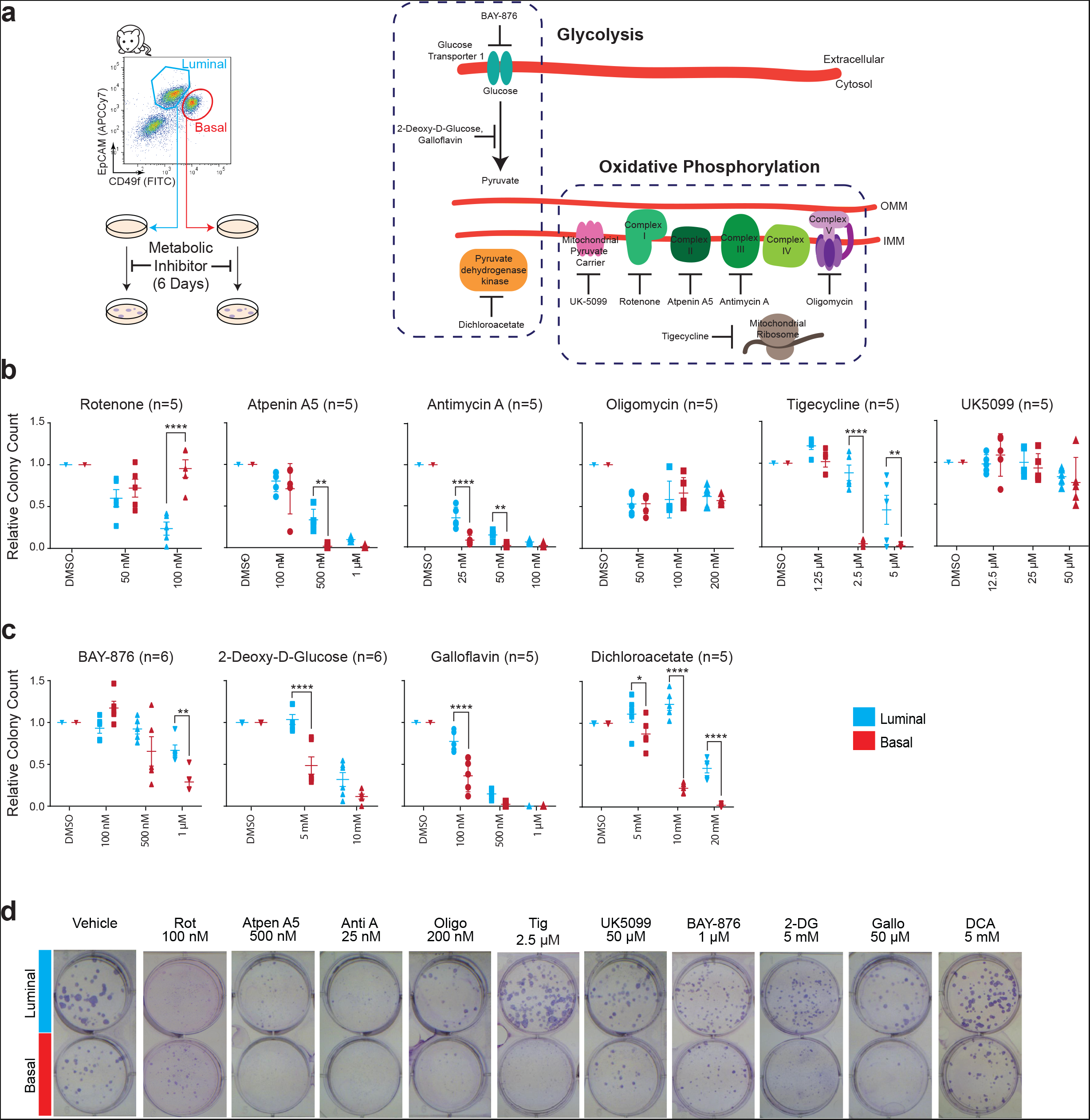
Metabolic inhibitors expose lineage-restricted vulnerabilities of MECs. A. FACS gating strategy and pictorial summary of metabolic inhibitors and their respective targets used to measure effects on mammary progenitor activity using the CFC assay. B. Dose-dependent effects of oxidative phosphorylation (OXPHOS) inhibitors on mouse mammary CFCs. Colony counts were normalized to their respective basal or luminal vehicle control;. Number of biological replicates per drug is shown in brackets. Matched-pairwise analysis. Data are mean ± SEM. * P≤0.05; ** P≤0.01; *** P≤0.001; ****P≤0.0001. C. CFC enumeration after treatment after treatment with glycolysis inhibitors D. Representative images of CFC plates after 6 days of culture, inhibitor concentration and colony type are indicated.

### Breast Cancer Subtypes Retain Metabolic Features of their Putative Cell-of-Origin

To interrogated whether any of the PAM50 breast cancer subtypes had significant enrichment of our MEC-specific metabolic network (Figure 4A) we performed single sample gene set enrichment analysis (ssGSEA)^42^ on breast cancer patients from the METABRIC database^43^. We observed striking relationships between the metabolic preferences of normal MEC and breast cancer subtypes (Figure 4A). Specifically, the highly mesenchymal Claudin-low subtype was most enriched for the basal cell metabolic network. The aggressive basal-like breast cancer was most significantly correlated to the luminal progenitor metabolic network (Figure 4A). Luminal A and B subtypes showed significant enrichment for the mature luminal metabolic network (Figure 4A). Thus, breast cancer subtypes displayed strong activity for metabolic cluster of their proposed cell-of-origin, suggesting that breast cancers co-opt the metabolic network of their precursor cells.

**Figure 4:**
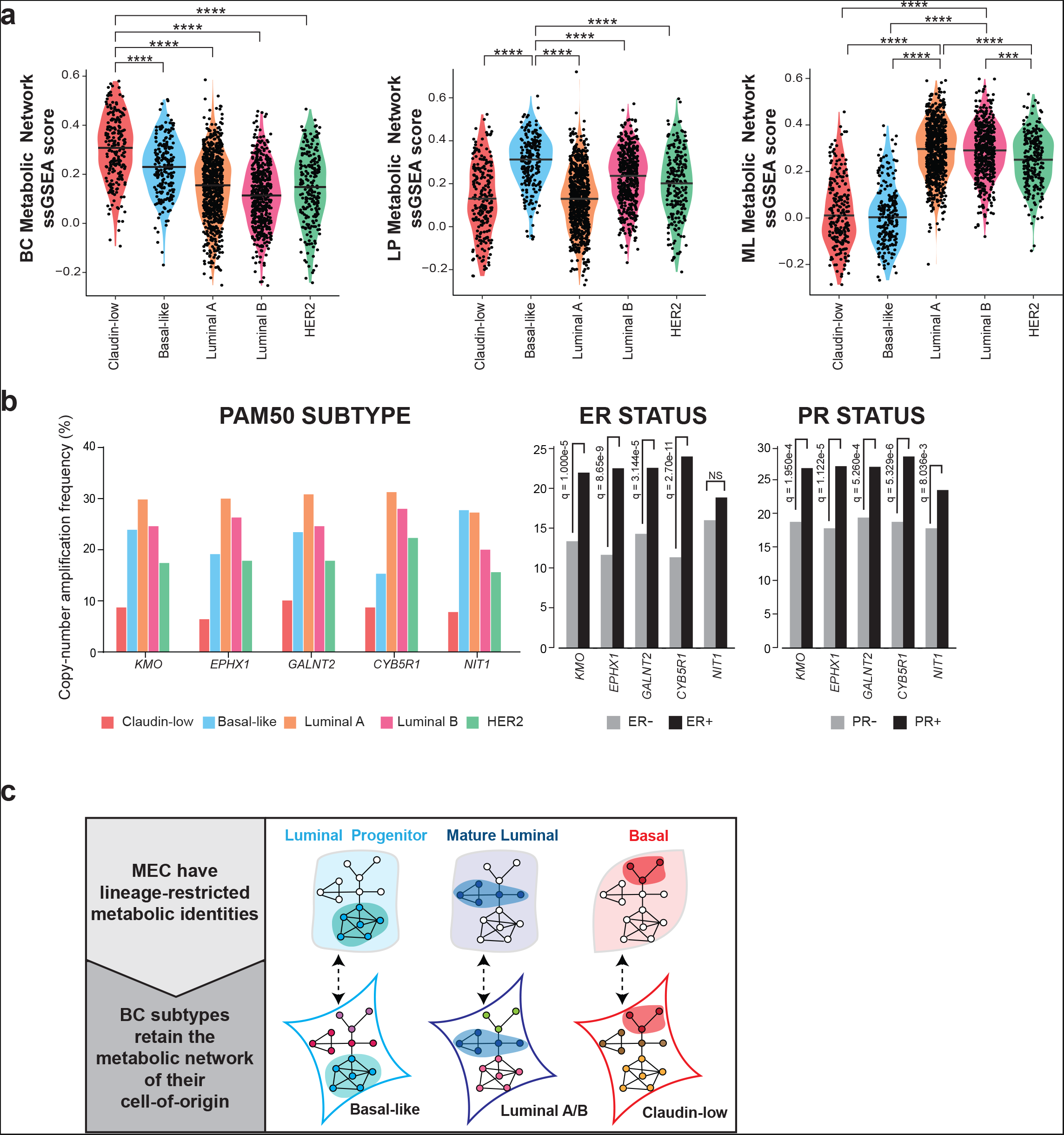
Breast cancer subtypes retain metabolic features of specific primary MECs. A. Violin plots of single sample gene set enrichment analysis (ssGSEA) scores comparing the metabolic network of basal, mature luminal and luminal progenitor cells to the PAM50 subtypes of breast cancer. Each dot represents a patient from the METABRIC study. One-way ANOVA in conjunction with a Tukey’s test was performed to determine the statistical significance of the differences in median ssGSEA scores for different breast cancer subtypes, all compared to the subtype with the highest median. *** P≤0.001; ****P≤0.0001. B. Copy-number amplification frequency of *KMO*, *EPHX1*, *GALNT2*, *CYB5R1* and *NIT* in the breast cancer patients from the METABRIC cohort ^43^, grouped based on PAM50 classifier, Estrogen receptor (ER) and Progesterone receptor (PR) status. KMO, EPHX1, GALNT2, CYB5R1 and NIT were found all highly abundant in the mature luminal metabolic network. C. Graphical abstract of the key findings in this study. Mammary epithelial cell types have lineage-restricted metabolic identities, as found by metabolic protein abundance, characterization of the mitochondria and drug effects on progenitor capacity. This is visualized by the distinct coloured nodes in each normal MEC population. The large amount of heterogeneity in breast cancer metabolism is represented by the unique colours for the nodes in each subtype. Part of this heterogeneity can be explained by the diverse cellular origin of breast cancer subtypes, where they inherit metabolic features of their cell-of-origin, projected here by the overlapping nodes with the same node colours as their primary MECs.

Recent studies have reported the successful targeting of metabolic vulnerabilities which are specific to the tissue-of-origin^16,44^ or stem from chromosomal abnormalities^45,46^. We therefore used cBioportal^47,48^ to determine copy number amplifications in metabolic genes within our MEC-specific networks in order to identify novel subtype-specific metabolic targets. Interrogation of our luminal progenitor metabolic network revealed *PHGDH*, a known amplified gene and selective vulnerability in ER- basal-like breast cancers^31,49^, the subtype thought to originate from luminal progenitors. We identified 5 other highly abundant proteins in our mature luminal metabolic network, namely EPHX1, NIT1, CYB5R1, GALNT2 and KMO, whose genes were amplified in ER+, PR+ as well as most consistently in Luminal A and B breast cancers (Figure 4B). These 5 proteins do not participate in the same metabolic pathway but are all found on chromosome 1q. Wholearm amplification of 1q together with 16q loss (+/−) is a hallmark chromosomal event in ER+ breast cancers^43,50,51^. The fact that we find metabolic network specific proteins being amplified at the chromosome level in the respective breast cancer subtypes points to these targets as possible cell-of-origin-specific metabolic vulnerabilities, which require further investigation.

## DISCUSSION

Using a combination of proteomics, characterization of the mitochondria and pharmacological inhibition, we uncovered distinct metabolic identities of the three normal mammary epithelial cell types (Figure 4C). This highlights a previously underappreciated metabolic heterogeneity present in the epithelial compartment of the normal human and mouse mammary gland. The observed lineage-drive metabolic programs may be intrinsic to cell identity or a reflection of cellular adaptations to distinct mammary microenvironments. Basal cells are in contact with the basement membrane which separates the epithelial layers from a complex mammary stroma composed of immune cells^52^, adipocytes^53^ and fibroblasts^54^. Coversely, luminal cells are exposed apically to the lumen of the mammary ductal tree. Since all our analyses were performed *ex vivo* on purified mammary cells, we reason that metabolic distinctions are hardwired and likely necessary to facilitate unique form and function of each mammary cell type. Our MEC-specific metabolic networks will enable further study into the influence of normal cells on the metabolic phenotype of known breast cancer subtypes. In addition, global proteomes of primary FACS-purified human and analogous mouse mammary cell types provide a valuable resource to further understand the regulatory networks that define these different epithelial lineages.

The metabolic phenotype of a cancer cell is dependent upon integrating multiple intrinsic and extrinsic cues^55^. The importance of the tissue-of-origin in tumor metabolism has now been established^13^. It has also been postulated that tumors located in the same tissue but derived from different cell(s)-of-origin would display different metabolic properties, however this has never been experimentally shown^56^. Our work demonstrates that part of the metabolic heterogeneity observed in breast cancers is instructed by the diverse cellular origins of these cancers (Figure 4C). For instance, Claudin-low, Basal-like, Luminal A & B appear to inherit metabolic features of basal cell, luminal progenitor and mature luminal populations, respectively. Arguably, cell lineage could be one of the most important determinants of cellular metabolism, as all perturbations (mutational or microenvironmental) will hijack the pre-existing metabolic network of the cell-of-origin as a backbone. Thus, in addition to mutational events, other characteristics of the tumor such as the cell-of-origin need to be considered in order to maximize the success of personalized cancer medicine. Our study lays the foundation for rationalized targeting of subtype-specific metabolic vulnerabilities, as informed by the metabolic networks of mammary epithelial cells.

## AUTHOR CONTRIBUTIONS

Conceptualization: MM, AEC, RK.

Methodology: MM, AEC, DP, CE, AS, VI, HK, VS.

Formal analysis: MM, AEC, KA, LP, MGV.

Resources: AS, TK, HB, CE, RK.

Writing: MM, TK, MAP, RK.

Visualization: MM, MA, KA.

Funding acquisition: TK, CE, RK.

Supervision: TK, MAP, RK.

## ACKNOWLEDGMENTS

We thank members of the Khokha Lab for reviewing this manuscript and helpful discussions. We are grateful to the Nanoscale Biomedical Imaging Facility at SickKids and the Princess Margaret and SickKids flow cytometry core facilities. This work is supported by funding from Canadian Institutes of Health Research (CIHR), Canadian Breast Cancer Foundation, and Canadian Cancer Society Research Institute. MM received a CIHR Masters Award.

## METHODS

### Human patient samples

All human tissue was acquired with patient consent and approval by the Institutional Research Ethics Board of the University of British Columbia (UBC; Vancouver, BC) and University Health Network (Toronto, ON). Hormonal status (premenopausal, follicular and luteal) was determined by a pathologist examining breast specimens at UBC (Ramakrishnan et al., 2002). Reduction mammoplasty specimens were minced and enzymatically dissociated in DMEM:F12 1:1 media with 15 mM HEPES plus 2% BSA, 1% penicillin-streptomycin, 5 μg/ml insulin, 300 U/ml collagenase (Sigma, C9891) and 100 U/ml hyaluronidase (Sigma, H3506) shaking gently at 37°C, overnight or for 16-18 hours. Epithelial organoids were harvested by centrifugation at 80g for 30 seconds and viably cryopreserved, as described previously (Labarge et al., 2013).

### Human breast single cell suspensions

Human breast tissue organoids were thawed and dissociated into single cell suspensions as reported previously (Eirew et al., 2010). Briefly, organoids were triturated in 0.25% trypsin-EDTA (Stem Cell Technologies, 07901) followed by 5 U/ml dispase (Stem Cell Technologies, 07913) and 50 μg/ml DNase I (Sigma, D4513) as described above for mouse samples, but for 5 minutes each. Cells were then washed in between steps with HBBS + 2% FBS and filtered using a 40 μm cell strainer.

### Human breast FACS staining

For FACS staining, antibodies against CD45 (PECy7), CD31 (PECy7), EpCAM (APCCy7) and CD49f (FITC) were used. Lineage (Lin) positive cells were defined as CD31^+^CD45^+^. Human mammary cell subpopulations were defined as: basal (Lin^−^ EpCAM^lo-med^CD49f^hi^); luminal progenitor (Lin^−^EpCAM^hi^CD49f^med^); mature luminal (Lin^−^ EpCAM^hi^CD49f^lo^). Dead cells were excluded following doublet exclusion using DAPI.

### Mice

All experiments were performed using 8-12 weeks old virgin female FVB wild-type mice (The Jackson Laboratory or Charles River). Mice were ovariectomized bilaterally, then allowed one week to recover. A slow-release 0.14 mg 17-β estradiol plus 14 mg progesterone pellet (Innovative Research of America) was then placed subcutaneously near the thoracic mammary gland for 2 weeks. This was done to obtain large quantities of viable mammary stem/progenitor cells for subsequent analysis, as previously reported (Casey et al., 2018; Shiah et al., 2015). All mice were cared for according to guidelines established by the Canadian Council for Animal Care under protocols approved by the Animal Care Committee of the Ontario Cancer Institute.

### Mouse mammary single cell suspensions

Harvested mammary glands were manually minced with scissors for 2 minutes, and then enzymatically dissociated using 750 U/ml collagenase and 250 U/ml hyaluronidase (Stem Cell Technologies, 07912) and diluted in DMEM:F12 for 1.5 hours. Samples were vortexed at the 1− and 1.5-hour mark. Red blood cells were lysed using ammonium chloride (Stem Cell Technologies, 07850). Cells were then mixed in trypsin-EDTA (0.25%, Stem Cell Technologies, 07901) that had been pre-warmed to 37°C using a 1mL pipette for 2 minutes. Next, they were washed in Hanks Balanced Salt Solution (HBSS) without calcium or magnesium plus 2% FBS and centrifuged at 350g. Finally, cells were mixed in dispase 5 U/ml (Stem Cell Technologies, 07913) plus 50 μg/ml DNase I (Sigma, D4513) for 2 minutes, washed in HBBS + 2% FBS and filtered using a 40 μm cell strainer to obtain single cells.

### Mouse mammary FACS staining

Dead cells were excluded following doublet exclusion using DAPI or Zombie UV Fixable Viability Kit (BioLegend) according to manufacturer’s instructions. For FACS staining, antibodies against TER119 (PECy7 or eFluor450), CD31 (PECy7 or eFluor450), CD45 (PECy7 or eFluor450), EpCAM (APCCy7), CD49f (FITC or PECy7), CD49b (PE) and Sca-1 (APC or Brilliant Violet 711) were used. Lineage (Lin) positive cells were defined as Ter119^+^CD31^+^CD45^+^. Mouse mammary cell subpopulations were defined as: total basal (Lin^−^ EpCAM^lo-med^CD49f^hi^); total luminal (Lin^−^EpCAM^hi^CD49f^lo^); luminal progenitor (Lin^−^ EpCAM^hi^CD49f^lo^CD49b^+^Sca-1^−^); mature luminal (Lin^−^EpCAM^hi^CD49f^lo^CD49b^−/+^Sca-1^−/+^). High and low mitochondrial activity populations were defined as MitoTracker Red^Hi^MitoTracker Green^hi^ and MitoTracker Red^lo^MitoTracker Green^hi^, respectively, and applied after gating for total luminal and basal populations. Fluorophores are specifically mentioned in individual figures. Cell sorting was performed on a BD FACSAria™ II.

### Mouse CFC assay

350 cells of the specified FACS-purified population were seeded together with 20,000 irradiated NIH 3T3 cells in a 6-well plate. Cells were cultured for 7 days at 5% oxygen in EpiCult-B mouse medium (Stem Cell Technologies, 05610) supplemented with 5% FBS, 10 ng/ml EGF, 20 ng/ml basic FGF, 4 μg/ml heparin, and 5 μM ROCK inhibitor (Millipore). Cells were allowed to adhere for 24 hours, and then either vehicle control (0.1% DMSO) or the indicated concentrations of inhibitors were added for the remaining six days.

### Mammary cell intracellular flow cytometry

All intracellular dyes were used to stain cells prior to cell surface marker staining protocol. Staining for total mitochondria (50 nM MitoTracker Green FM, Thermo Fisher, M7514), mitochondrial activity (250 nM MitoTracker Red CMXRos, Thermo Fisher, M7513), mitochondrial ROS (5 μM MitoSOX, Thermo Fisher, M36008), and cytosolic ROS (5 μM CellROX Green, Thermo Fisher, C10492) was performed by incubating cells at 37°C for 20-30 minutes following the manufacturer’s protocols and directly analysed without fixing. Cell analysis was performed in BD Biosciences Fortessa. Median fluorescent intensity (MFI) refers to the fluorescence intensity of each event (on average) of the selected cell population, in the chosen fluorescence channel (PE Texas Red or FITC) and was determined by using the flow cytometry analysis software FlowJo.

### Metabolic inhibitors used *in vitro*

Vehicle and drugs were added such that the final concentration of DMSO did not exceed 0.1% (vol/vol). The following drugs were used in this study: 2-Deoxy-D-glucose (Sigma; D8375), dichloroacetate (Sigma; 347795), BAY-876 (Structural Genomics Consortium), rotenone (Sigma; R8875), tigecycline (CarboSynth, 220620-09-7), antimycin A (Sigma, A8674), oligomycin (Sigma, 75351), atpenin A5 (Cayman Chemicals, 11898), UK-5099 (Sigma, PZ0160), galloflavin (Sigma; SML0776).

### Transmission electron microscopy

Mammary epithelial cells were FACS-purified from 3 EP-treated ovariectomized 8-12 week old mice. Cells were pooled together to increase yield and then pelleted for 5 mins at 4°C at max speed. Supernatant was removed and then fixed with 2% glutaraldehyde in 0.1 M sodium cacodylate buffer pH 7.3, without disturbing the pellet. Samples were processed by the Nanoscale Biomedical Imaging Facility (SickKids, Toronto, ON). Images were acquired using the FEI Technai 20 transmission electron microscope. Scale bars are specific to images.

### Seahorse

MEC subpopulations (Luminal progenitor, mature luminal and basal cell) were FACS-purified from unstaged mice and 10,000 cells were plated into each well of collagen pre-coated Seahorse plates. The cells were culutred in the 5% O2 incubator for 6 days to reach at least 80-90% confluence. On the 7th days, cells were switched to DMEM:HAM’s F12 with no bicarbonate containing 5% FBS, insulin (Thermo Fisher, 12585014), EGF (STEMCELL Technologies; 78006.1), bFGF (STEMCELL Technologies), hydrocortisone (STEMCELL Technologies, 78003.1), Rock inhibitor (Millipore, SCM075) in 5% oxygen conditions. Then the plate was allowed to equilibrate for 1 hour in the Seahorse incubator. Inhibitors used for the assay include oligomycin (2 μM), FCCP (1 μM, Sigma, C2920) and antimycin A (1 μM). After the assay, cell viability was determined using the CyQUANT nuclear dye (Thermo Fisher, C35007). Data was analyzed on the WAVE platform and normalized to the number of live cells determined after the viability assay.

### Proteomics on FACS-purified human mammary subpopulations

For Liquid Chromatography-Mass Spectrometry (LC-MS) of human mammary subpopulations, 100,000 cells from each population were isolated from each patient, as described (Casey et al., 2018). After FACS purification, cells were washed in ice-cold PBS and pelleted. Pellets were then resuspended in 50% (vol/vol) 2, 2, 2-trifluoroethanol in PBS and disrupted into cellular lysates sequentially by repeated probe sonication, followed by six freeze-thaw cycles. Proteins in cellular lysates were denatured by incubation at 60°C for 2 h, oxidized cysteines reduced using 5 mM dithiothreitol for 30 min at 60°C and alkylated through reaction with 25 mM iodoacetamide for 30 min at room temperature in the dark. Each sample was diluted five times using 100 mM ammonium bicarbonate, pH 8.0. Proteins were digested into peptides through addition of 5 μg of MS-grade trypsin (Promega). The digestion was performed overnight at 37°C and subsequently desalted using OMIX C18 pipette tips (Agilent). Peptides were semidried through vacuum centrifugation and resuspended in water with 0.1% formic acid. Subsequently, all samples were analyzed using an Easy-LC1000 (Thermo Fisher Scientific) coupled to the Orbitrap Fusion tandem mass spectrometer (Thermo Fisher Scientific). Peptides were separated on an ES803 (Thermo Fisher Scientific) nano-flow column heated to 50°C using a 4-h reverse-phase gradient.

### Bioinformatics Analysis of human mammary subpopulation proteomes

#### Proteomics Processing

Mass spectrometric data was analyzed using the MaxQuant quantitative proteomics software (version 1.5.8.3) and a Human UniProt sequences FASTA database (complete human proteome; release 2015-01, 42,041 sequences). Carbamidomethylation of cysteine was specified as a fixed modification and oxidation of methionine was specified as a variable modification. Proteins were identified with a minimum of two razor+unique peptides, the maximum false peptide discovery rate was specified as 1%, and “match between runs” was enabled. The distribution of intensity-based absolute quantification (iBAQ) values was adjusted to the distribution of label-free quantification (LFQ) values based on the median for each sample. This allowed for imputation of missing LFQ values with iBAQ values(Wojtowicz et al., 2016). Non-zero values were log2-transformed. The final list consisted of 6034 unique protein groups detected in at least one of the samples. Further data processing was performed using the R statistical environment (version 3.5.2) (Bunn and Korpela). For protein groups in which both LFQ and iBAQ values were missing, the 0 values were imputed with a random value between 1 and 1.5. Imputation was performed as a precautionary measure for further statistical analysis. As four samples were run on a separate day, intensity values were then adjusted for sample batch effects using the ComBat method in the surrogate variable analysis “sva” R package (version 3.30.1) (Johnson et al., 2007; Leek et al.).

#### Total Proteome Bioinformatics

Non-imputed ComBat-modified iBAQ-adjusted LFQ values were used to discover uniquely expressed proteins in each cell type. Averages across samples in each cell type were taken, resulting in one mean expression value for each protein in each cell type (*n*_BC_ = 9, *n*_LP_ = 10, *n*_ML_ = 10). Next, the values of zero for each cell type and associated proteins were excluded from the analysis. Number of proteins expressed in each cell type were summarized in a Venn diagram, created using the “VennDiagram” R package (version 1.6.20) (Chen and Boutros, 2011). Gene set enrichment was conducted on the same values. Mean expression values for each protein in each of the cell types were ranked according to descending log2 median intensities and grouped into deciles. The protein with the highest intensity received a rank of 1 and thus, was placed in the first decile. Meanwhile, the protein with the lowest mean intensity received a rank of *y* and was placed in the tenth decile, where *y* represents the total number of proteins detected in a particular cell type. Pathway analysis via the “enrichR” R package (version 1.0) was conducted on the proteins in each decile (Chen et al., 2013; Kuleshov et al., 2016).

Principal component analysis (PCA) was performed by calculating Euclidean distances of scaled expression values. PCA scores were plotted in a plane defined by the first two components (that is, PC1 and PC2) using the “ggbiplot” R package (Vu, 2019). Ellipses were drawn around cell type clusters, where centroids were the barycentre of each cluster and the diameter represented the maximum variance.

Heat maps depicted z-scores of protein expression values (*x*) computed using the formula: (*x* – mean(*x*))/standard_deviation(*x*). Divisive hierarchal clustering dendograms of Pearson distance matrices for samples and proteins were created using DIANA (DIvisive ANAlysis Clustering) method in the “cluster” R package (version 2.0.7-1) (Maechler et al.). Heat maps were plotted using the “pheatmap” (version 1.0.12) and “RColorBrewer” (version 1.1-2) R packages (Kolde, 2019; Neuwirth, 2014).

#### Metabolic Cluster Derivation and Pathway Analysis

A metabolic proteome was obtained by filtering the total proteome using a curated list of 2753 genes that encompasses all known human metabolic enzymes and transporters (Possemato et al., 2011). Based on matching by gene symbols, 1020 proteins related to metabolism were found in the total proteome of 6034 proteins, including “PKM” which was not identified in the curated list. As multiple protein groups in the proteome shared the same gene symbols, duplicates were included in the analysis. Metabolic signatures were acquired by looking at proteins in which mean expression met the fold-change and statistical change cut-offs in each cell type compared to the other two cell types (*n*_BC_ = 9, *n*_LP_ = 10, *n*_ML_ = 10). The log2 fold-change (FC) cut-off was greater than 0 and the statistical significance cut-off was P < 0.05 in a one-way ANOVA and Tukey’s multiple comparisons test. Pathway analysis of metabolic clusters was conducted using Enrichr (https://amp.pharm.mssm.edu/Enrichr/). Enrichr is a comprehensive gene set enrichment tool that is available both as a web interface (Chen et al., 2013) and an R package (Kuleshov et al., 2016). It queries a list of gene symbols and returns commonly annotated pathways by searching large gene set libraries. The gene set library selected for our analysis was Gene Ontology Biological Process (GOBP) 2018. For each cell-type signature, the top ten GOBP terms enriched by gene sets were sorted by lowest to highest combined score (ln(p-value) *z-score), a metric used by Enrichr to find the best ranking terms compared to other methods.

#### Correlations to PAM50 Breast Cancer Subtypes

Gene expression for PAM50 breast cancer subtypes (Her2, Luminal A, Luminal B, Basal-like, and Claudin-Low) and clinical annotations was performed in the METABRIC cohort (Curtis et al., 2012) and was obtained from cBioPortal (Cerami et al., 2012; Gao et al., 2013). It provided gene expression profiles and classified breast cancer subtypes for 1980 patients. The gene expression profiles for the breast cancer subtypes (were correlated to our metabolic signatures via single-sample Gene Set Expression Analysis (ssGSEA) using the “GSVA” R package (version 1.30.0) (Hänzelmann et al., 2013). ssGSEA scores for each signature in the breast cancer subtypes were assessed for significance using a one-way ANOVA and student’s t-test.

#### Statistical Analysis and Reproducibility

All details pertaining to biological “n” numbers or error bars can be found in the relevant figure legends. Details pertaining to the statistical analysis of global and metabolic proteome can be found in the relevant methods section detailing bioinformatics analyses. Statistically significant differences are indicated by asterisks, which denote size of significance levels (p-values: ns P > 0.05; * P ≤ 0.05; ** P ≤ 0.01; *** P ≤ 0.001; **** P ≤ 0.0001.) For intracellular flow cytometry analysis statistical significance was calculated using two-way ANOVA and Tukey’s multiple comparisons test. For *in vitro* clonogenic assays comparing high and low mitochondrial mammary cells, a two-way ANOVA and Bonferonni’s multiple comparison test was used. For *in vitro* clonogenic assays, statistical significance for all drug testing comparisons was calculated using two-way ANOVA and Sidak’s multiple comparisons test.

#### Data Availability

The mass spectrometry data associated with this manuscript will be submitted to a public repository (the Mass spectrometry Interactive Virtual Environment; http://massive.ucsd.edu). These data are associated with the identifier at FTP download site:. The mouse mammary proteome data (used in Figure 2E, S3B) is published (Casey et al., 2018) and can be downloaded from the FTP download site: ftp://MSV000079330@massive.ucsd.edu with the identifier MSV000079330

---

**Figure S1:**
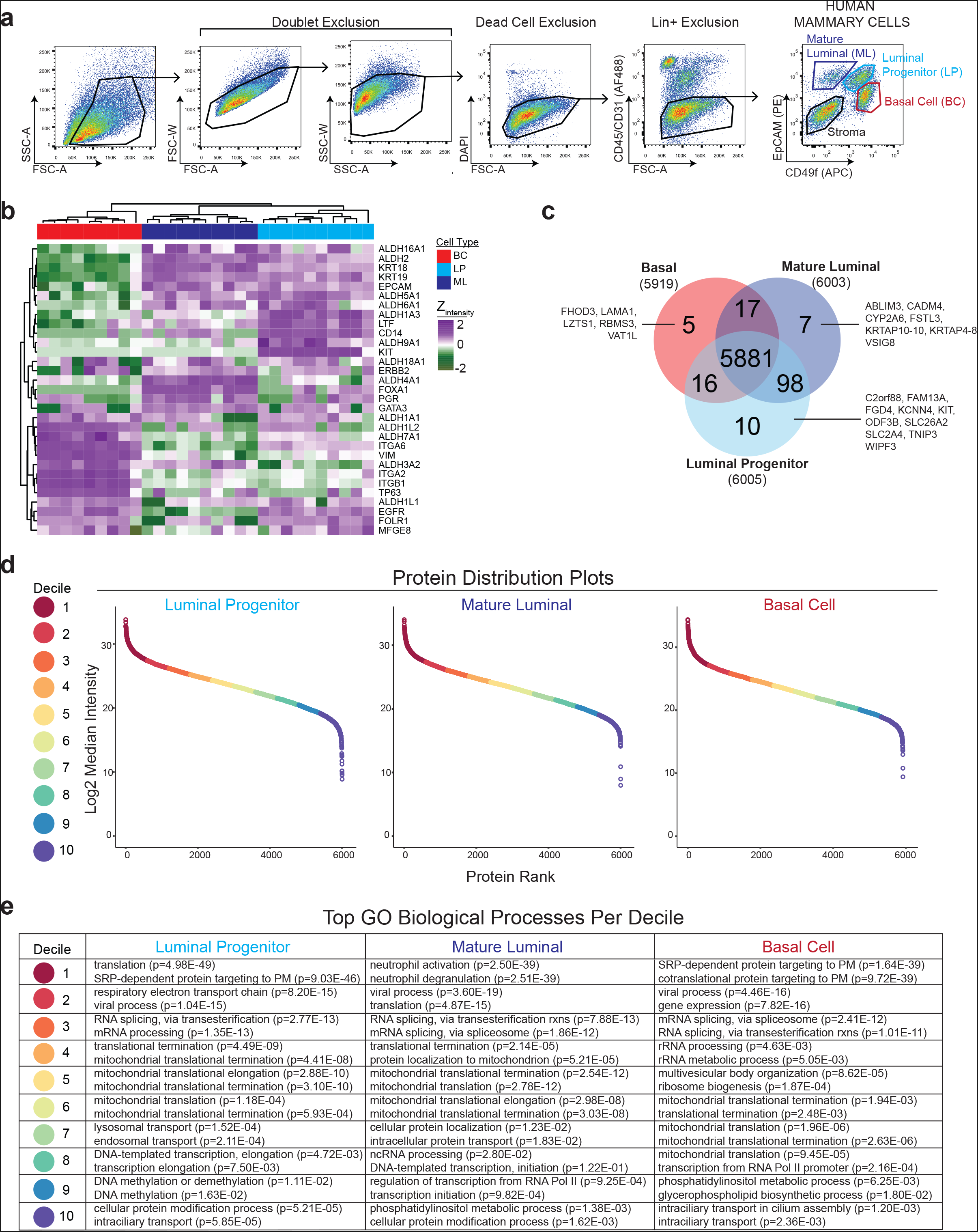
Characterization of proteomic datasets of primary FACS-purified human mammary epithelial cells. A. Gating strategy for FACS-purifying human mammary epithelial cells. Total cells from dissected human breast tissue are gated to exclude debris. Doublet, dead cell and Lineage (Lin+) exclusion ensures sorting of single, live and non-immune cells. B. Heatmap shows unsupervised hierarchical clustering and abundance of a set of known marker proteins well established for distinguishing mammary epithelial cell types. C. Venn diagram summarizing the distribution of the 6040 detected proteins among mammary populations. The numbers in brackets are the total number of proteins detected in that cell type. D. Pathway analysis using Enrichr was performed on each decile for each MEC type. The top 2 GO Biological Processes per each decile are summarized with its associated adjusted p-value in brackets.

**Figure S2:**
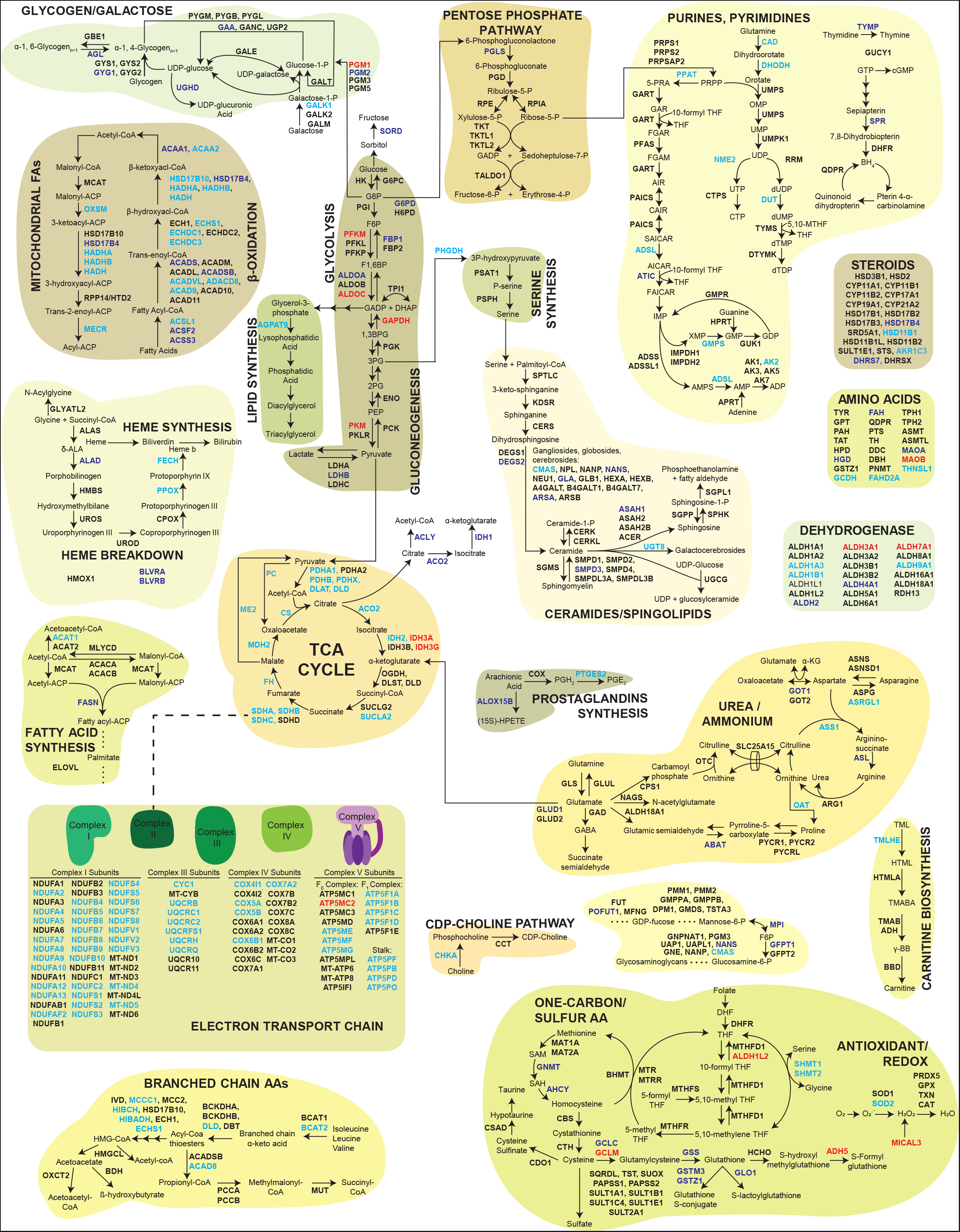
Map of human mammary epithelial cell metabolism. A. Metabolic map is adapted from a previously published template ^34^. Proteins are coloured-coded to denote which mammary cell-type specific metabolic network they demonstrated their highest expression level (Black = not significant or not detected, Light blue = luminal progenitors, Dark blue = mature luminal and red = basal).

**Figure S3:**
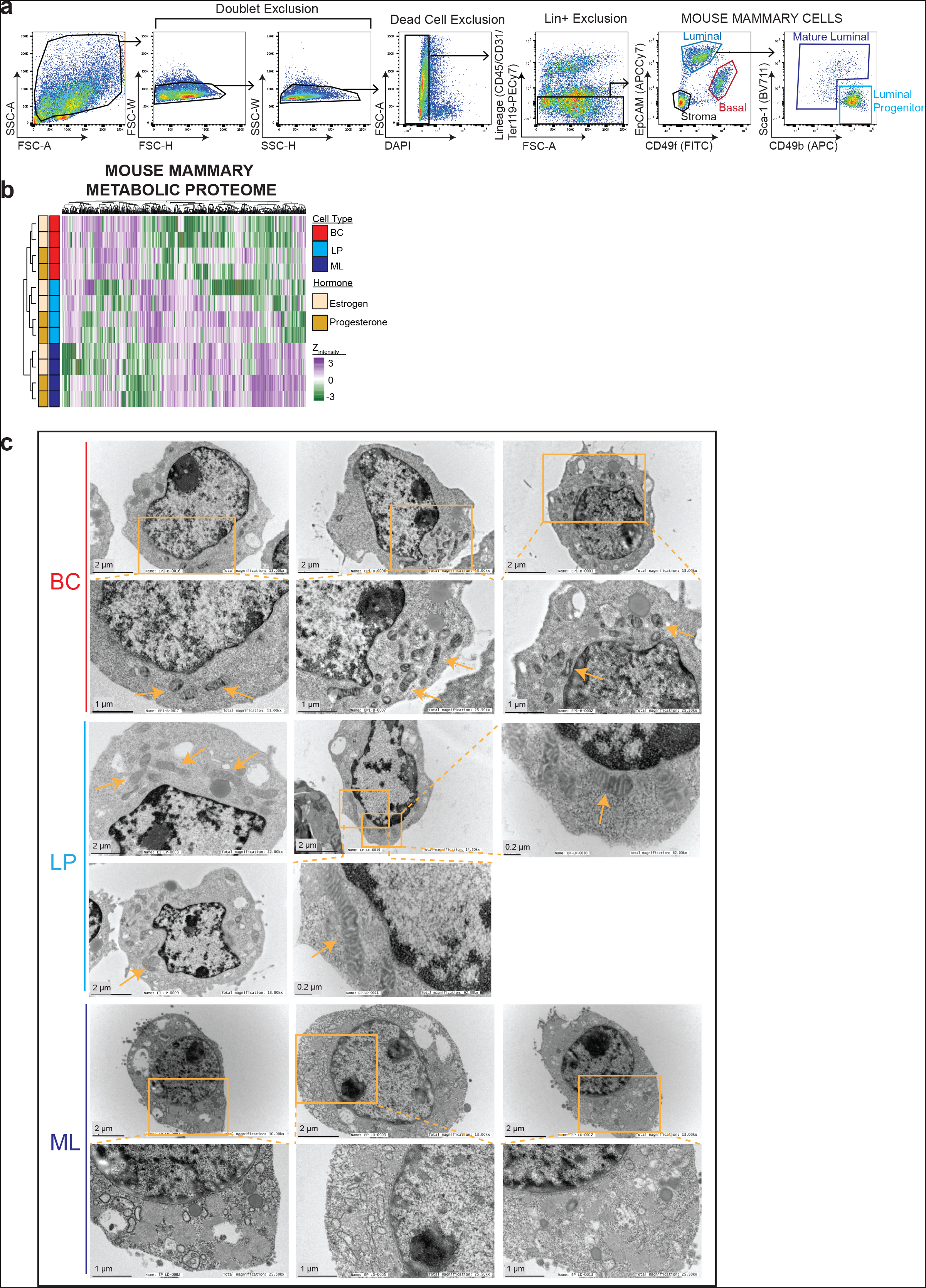
Characterization of the mouse mammary mitochondria. A. Gating strategy for FACS-purifying mouse mammary epithelial cells. Total cells from dissected mouse mammary gland are gated to exclude debris. Doublet, dead cell and Lineage (Lin+) exclusion ensures single, live and non-immune cells are analyzed. B. Heatmap showing unsupervised hierarchical clustering and z-scores of only the metabolic proteins, determined by a curated list ^31^, from our previously published mouse mammary proteomic dataset ^25^. C. Representative transmission electron micrographs of FACS-sorted mammary cell pellets. Arrows indicate mitochondria. Magnifications are specified in each image.

**Figure S4:**
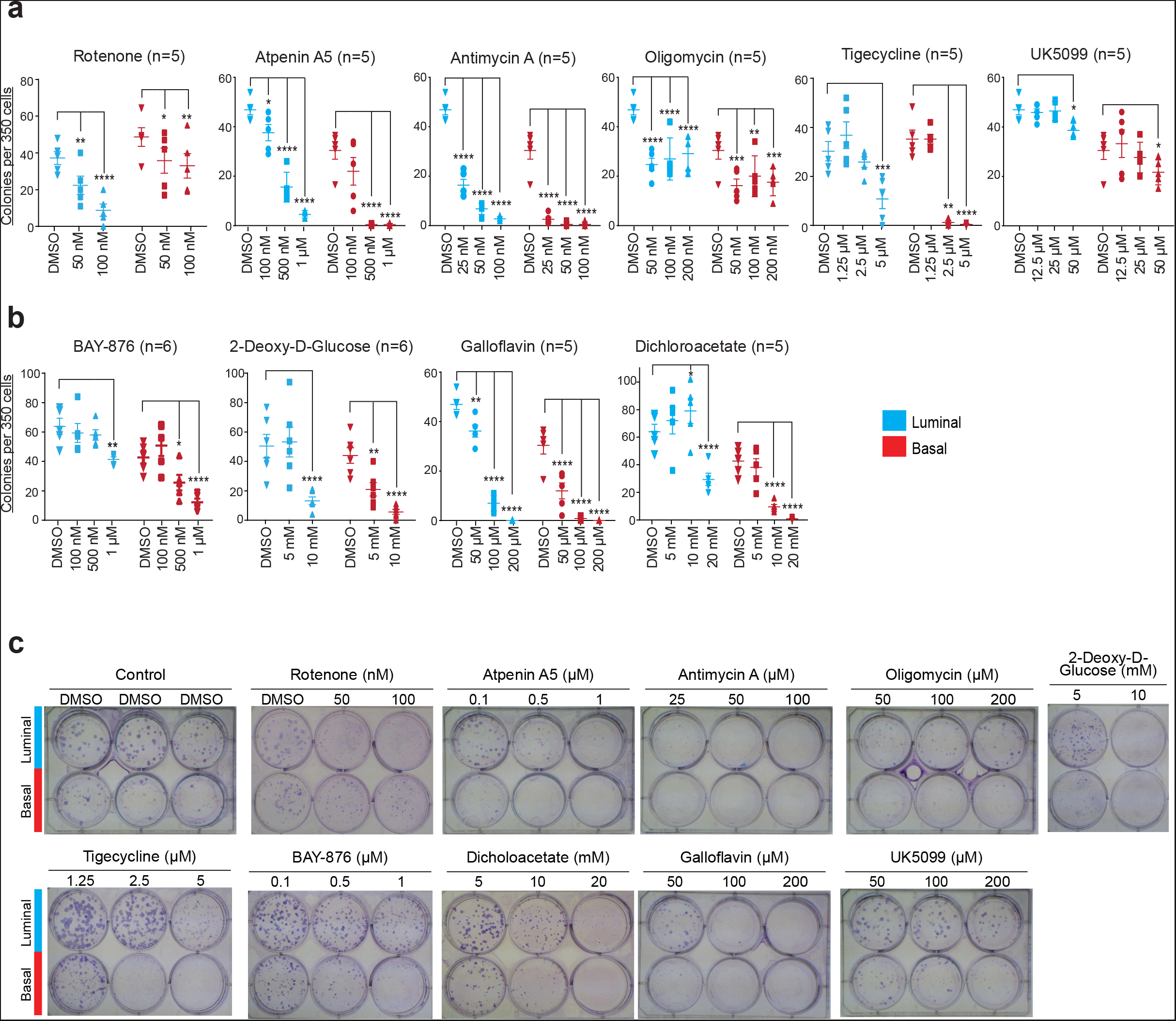
Absolute CFCs of mammary lineages following treatment with metabolic inhibitors. A. Quantification of absolute CFC counts at various concentrations of the specified OXPHOS inhibitor. Basal colonies are red and luminal colonies are blue. Each dot represents a mouse and number of biological replicates per drug is shown in brackets. Data are mean ± SEM. * P≤0.05; ** P≤0.01; *** P≤0.001; ****P≤0.0001. B. Quantification of absolute CFCs after treatment with glycolysis inhibitors. C. Representative images of CFC plates at the end of 6-day treatment. Inhibitors, concentrations and the mammary cell types are indicated.

